# What Do Biological Foundation Models Compute? Sparse Autoencoders from Feature Recovery to Mechanistic Interpretability

**DOI:** 10.64898/2026.03.04.709491

**Authors:** Alexey V. Orlov, Yulia V. Makus, German A. Ashniev, Natalia N. Orlova, Petr I. Nikitin

## Abstract

Foundation models trained on protein and DNA sequences are increasingly deployed for variant interpretation, drug design, and gene regulation prediction, yet their internal representations remain opaque – limiting both biological insight and trust in model-guided decisions. Existing interpretation approaches establish what these models encode but cannot reveal how biological knowledge is internally organized and computed. Sparse autoencoders (SAEs) offer a complementary approach by decomposing model activations into interpretable features, each capturing a distinct biological concept. Over the past year, SAEs have been applied to protein language models, genomic language models, pathology vision transformers, single-cell foundation models, and protein structure generators. Here we provide a systematic review of sparse dictionary learning across biological foundation models. We find that independent studies using different architectures and evaluation strategies consistently recover features spanning biological scales – from secondary structure elements and functional domains in proteins to transcription factor binding sites and regulatory elements in genomes – providing convergent evidence that these models learn interpretable representations accessible through sparse decomposition. However, we identify a critical gap: validation relies almost exclusively on matching features against existing annotations, risking circularity when those annotations derive from the same sequence databases used for model training. We propose a three-level interpretability framework – representational, computational, and causal mechanistic – and argue that the field’s most distinctive opportunity lies in experimental validation through deep mutational scanning, massively parallel reporter assays, and structural characterization, which can establish whether these models have learned genuine biological mechanisms rather than training set statistics.

## 1. Introduction

Foundation models have transformed computational biology [1,2]. Protein language models trained on evolutionary sequence data achieve state-of-the-art performance across structure prediction, function annotation, and variant effect estimation; genomic language models pretrained on multi-species DNA capture regulatory logic, gene expression prediction, and chromatin accessibility without task-specific supervision [3–7]. These successes raise a natural question: what biological knowledge do these models encode, and how is it organized? Understanding how structural and functional information emerges from sequence data alone may illuminate principles of biological information encoding; deploying these models for drug design, variant interpretation, or synthetic biology requires confidence that predictions reflect genuine biological mechanisms rather than dataset artifacts.

Two generations of interpretability methods have addressed this question with fundamentally different objectives. Behavioral methods – attention analysis, attribution, probing, and Jacobian-based dependency analysis – characterize what information is encoded in model representations and which inputs drive predictions [8–11]. These approaches have established that protein language models encode residue-residue contacts, secondary structure, and functional annotations, while genomic models capture regulatory elements, splice site logic, and RNA secondary structure [4–6,8,12]. However, behavioral methods share a critical limitation: they describe model outputs without revealing how information is organized and computed internally [9,13].

Sparse autoencoders (SAEs) offer a qualitatively different approach. Rather than querying the model from outside, SAEs decompose high-dimensional activation spaces into overcomplete dictionaries of sparsely activating features, aiming to identify the fundamental units of model computation [14,15]. The methodology originated in the mechanistic interpretability of large language models [16,17] and has been rapidly transferred to biological foundation models since late 2024. Over a period of less than one year, SAE analyses have been applied to protein [18–21] and genomic [22–24] language models of varying scale, pathology vision transformers [25], single-cell foundation models [26], and protein structure generators [27], recovering features that span biological scales from individual nucleotide identity to multi-domain protein architecture. The term “autoencoder” has extensive prior use in computational biology – variational and denoising autoencoders are widely employed for single-cell RNA sequencing [28] – but these are models of data that learn to reconstruct expression profiles, whereas interpretability SAEs are tools for decomposing internal activations of already-trained models into interpretable components.

Despite this rapid progress, no systematic review has examined sparse dictionary learning across biological foundation models. Existing reviews of foundation models in bioinformatics address model architectures and benchmark performance but treat interpretability peripherally: Li et al. [29] and Guo et al. [30] survey model families without analyzing how internal representations are organized, and Liu et al. [31] cover biomedical foundation models broadly without dedicated discussion of mechanistic interpretability methods. From the interpretability side, Shu et al. [32] provide a comprehensive survey of SAEs for large language models but do not address biological applications – where data structure, validation opportunities, and downstream utility differ fundamentally from natural language. This review bridges these two bodies of work. Because the field has developed almost entirely since late 2024, a substantial fraction of the primary literature exists only as preprints; we include these to provide a comprehensive account of current progress while noting that peer-reviewed validation of many findings is still pending. Our central argument is that sparse dictionary learning enables a shift from characterizing what models encode to investigating how biological knowledge is internally organized, but that a significant distance remains between recovering interpretable features and establishing mechanistic understanding. Bridging this distance requires validation beyond annotation matching – causal interventions and experimental verification. We develop this argument by first surveying behavioral interpretation methods to delineate the explanatory limits that motivate sparse dictionary learning, then tracing SAE architectural evolution and its mapping to biological objectives, synthesizing applications across protein, genomic, and alternative modalities, and concluding with open questions that determine whether SAE-based interpretability can advance beyond feature cataloging toward genuine biological understanding.

## 2. Behavioral Interpretation Methods: Achievements and Explanatory Limits

Behavioral interpretation methods analyze trained models without modifying their parameters, establishing what biological knowledge foundation models have acquired and how they process sequence information. Four approaches have been widely applied to protein and genomic language models: attention analysis, attribution methods, probing tasks, and Jacobian-based pairwise dependency analysis (Figure 1). Each addresses a distinct analytical question and has produced substantial biological findings. This section surveys their contributions with particular emphasis on the explanatory boundaries that, collectively, motivate the transition to sparse dictionary learning.

**Figure 1.**
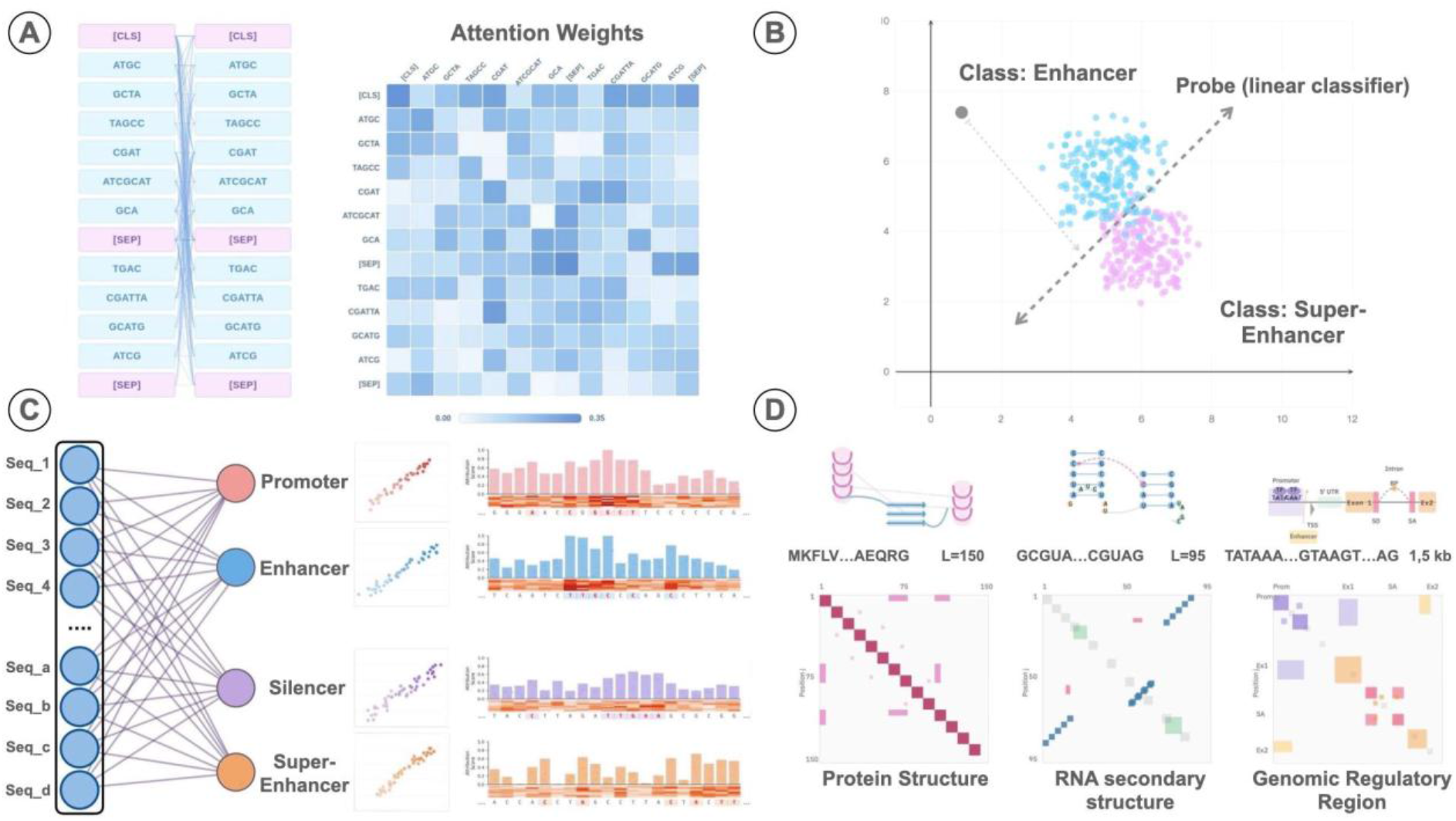
Overview of behavioral interpretation methods for biological foundation models. (A) Attention analysis extracts pairwise attention weights between sequence tokens, revealing positions the model considers related. (B) Probing tasks train linear classifiers on frozen embeddings to assess what biological properties are decodable from representations. (C) Attribution methods assign importance scores to input features, quantifying their contribution to model predictions. (D) Jacobian-based pairwise dependency analysis measures how perturbations at one position affect predictions at others, yielding dependency maps that capture structural relationships in proteins, RNA secondary structure, and genomic regulatory regions. Adapted from [33–36].

### 2.1 Attention Analysis

Transformer architectures compute pairwise attention between input tokens, and these weights remain accessible after training (Figure 1A). Systematic analysis of attention in protein language models revealed correlations with known structural features. Rao et al. showed that attention heads in ESM capture residue-residue contacts, with a sparse logistic regression on attention maps achieving long-range Top-L precision of 41.1% on 14,842 test structures – outperforming unsupervised coevolution-based methods [37]. Probing of frozen ESM representations separately confirmed that secondary structure, tertiary contacts, and binding sites are encoded in the learned embeddings [6]. Layer-wise analysis of attention demonstrated that deeper layers encode progressively more complex biophysical properties [10]. These patterns proved consistent across architectures, with MSA Transformer exhibiting distinct row-attention for structural contacts and column-attention for positional covariation across homologous sequences [38,39].

Analogous patterns emerged in genomic language models. After fine-tuning on respective tasks, DNABERT attention concentrates on transcription factor binding sites, TATA boxes, and intronic regions flanking splice sites [40]. Nucleotide Transformer models from 500M to 2.5B parameters show attention head specialization for enhancers, promoters, and UTR regions, with the Multispecies 2.5B model showing 117 of 640 heads enriched for introns and the 1000G 2.5B model achieving nearly 100% attention on enhancer regions – patterns emerging without task-specific supervision [4]. SpliceBERT exhibits elevated attention between donor and acceptor sites within the same intron, reflecting mechanistic coupling of splice site recognition [41].

However, the interpretive value of attention weights is contested. Jain and Wallace demonstrated that adversarially constructed attention distributions with near-maximal divergence from learned weights yield negligible change in model output, establishing that attention and prediction can be substantially decoupled [42]. Subsequent work clarified conditions under which attention can be informative [43,44], but identified further complications: information mixing across layers undermines single-layer interpretation [45], and attention weights capture only part of token influence, as transformed vector norms also contribute to final representations [46]. Notably, the original DNABERT study itself relied on attention visualization as its primary interpretability tool for identifying transcription factor binding sites and promoter motifs [40], yet the limitations above suggest that such attention-based biological interpretation should be treated with caution. Attention analysis thus reveals what positions the model considers related, but neither whether these relationships drive predictions nor how they are computationally implemented.

### 2.2 Attribution Methods

Where attention reveals positional relationships, attribution methods quantify which inputs drive specific predictions by assigning importance scores to individual positions (Figure 1C). The methodological foundations include Integrated Gradients [11], SHAP [47], DeepLIFT [48], and in-silico saturation mutagenesis (ISM) [49]. These methods were extensively applied to supervised genomic models [50,51] before being adopted for foundation models.

For protein language models, ESM-1v demonstrated that masked marginal probability differences enable zero-shot variant effect prediction across 41 deep mutational scanning datasets [52]. Integrated Gradients applied to protein models identified functionally important residues at the RNA binding interface of Pab1, consistent with known structure-function relationships [53]. Genomic foundation model publications rely predominantly on attention visualization; gradient-based attribution appears more frequently in downstream applications, as in GENA-LM’s use of ISM to demonstrate sensitivity to splice site substitutions [54].

Applying gradient-based methods to biological sequences presents domain-specific challenges beyond the general limitations known from general deep learning and computer vision [55,56]. Majdandzic et al. identified a fundamental issue: biological sequences are encoded as one-hot vectors constrained to a simplex, but networks learn functions across the full embedding space, including regions no valid sequence can occupy [57]. Gradients pointing outside the simplex do not correspond to meaningful sequence changes, yet standard methods report them as if they did – an artifact affecting 10–20% of positions [57]. Additional considerations include baseline selection, where dinucleotide-shuffled references outperform zero baselines [58], and integration path choice [59]. Computational cost constrains ISM for longer sequences, but two algorithmic innovations have made exhaustive mutagenesis tractable for multi-kilobase genomic contexts: fastISM exploits shared computation between wild-type and mutant forward passes, achieving 10× speedup for models with wide early layers [60], while compressed sensing ISM approximates the full mutation matrix from a small number of random perturbations, reducing compute by an order of magnitude with minimal accuracy loss [61].

Attribution methods thus identify which positions matter for individual predictions, but – like attention – they operate on single inputs without assessing what biological knowledge the model has acquired across its training distribution. This broader question is explored in particular in the probing tasks approach.

### 2.3 Probing Tasks

Rather than explaining individual predictions, probing asks what information is encoded in model representations across the entire sequence space. The approach involves training a simple classifier – typically linear – on frozen embeddings to predict biological properties (Figure 1B). If the classifier succeeds, the property is considered decodable from the representations [62,63].

Systematic benchmarking of protein language models established that self-supervised pretraining captures substantial structural and functional information [64]. Scaling to 650 M parameters, ESM-1b achieved Q8 secondary structure accuracy of 71.6% on CB513, competitive with HMM sequence profiles, and top-L long-range contact precision of 35.9% on CASP13 free-modelling domains [6]. ProtT5-XL, trained on up to 393 billion amino acids, the best-performing variant, reached Q3 of 87% for secondary structure and Q2 of 91% for membrane classification [5]. Layer-wise studies reveal task-dependent variation: different layers encode distinct biological properties, and the optimal layer rarely corresponds to the final one [18,19].

Genomic language models show parallel patterns. On the GUE benchmark [65], Nucleotide Transformer achieves MCC of 96% for splice sites and AUC of 0.92 for DNase I hypersensitive sites [4]. The BEND benchmark, extending evaluation to sequences up to 100kb, revealed that DNA models approach expert methods but weakly capture long-range dependencies [66]. HyenaDNA achieves state-of-the-art on 12 of 18 benchmark datasets with 1500-fold fewer parameters than Nucleotide Transformer, suggesting that architectural efficiency rather than scale may drive performance on certain tasks [67]. As with protein models, the optimal probing layer is task-dependent – for histone modification prediction, the gap between optimal and final layer reaches 15% relative performance [4].

These quantitative successes require careful methodological interpretation. Hewitt and Liang demonstrated that MLP probes achieve high accuracy even on random control tasks, indicating that probe complexity can confound results [68]. Minimum description length probing addresses this by measuring compression rather than raw accuracy [69]. More fundamentally, Belinkov formalized the distinction between information encoded in representations and information actually used by the model during inference [9]. Amnesic probing demonstrated that removing a property from representations may minimally affect downstream performance despite high probe accuracy – probing establishes correlation, not causation [70].

Critical evaluations specific to biological models have tempered initial optimism. Tang and Koo found that genomic language models consistently underperform supervised models on regulatory genomics tasks, arguing that existing benchmarks use metrics disconnected from biological utility [71]. Evo 2 underperformed relative to one-hot encoding for modeling non-coding regulatory logic [72], while for single-cell perturbation prediction, mean-of-training-examples baselines outperformed scGPT and scFoundation [73]. These findings underscore that high probe accuracy on standard benchmarks does not guarantee biological utility – current foundation models may capture shallow statistics rather than mechanistic logic, particularly for regulatory sequence analysis.

Probing reveals what information is linearly decodable from representations, but tests for predefined properties in isolation without capturing the relational structure inherent to biological sequences – the pairwise dependencies that underlie protein folding, RNA structure, and regulatory interactions.

### 2.4 Jacobian-based Pairwise Dependency Analysis

Biological sequences are fundamentally characterized by pairwise relationships: residue-residue contacts determine protein structure, base pairing defines RNA secondary structure, and regulatory elements interact across genomic distances. Jacobian-based analysis addresses this relational structure by measuring how perturbations at one position affect predictions at others, yielding dependency maps directly comparable to classical coevolutionary metrics (Figure 1D).

The conceptual foundation lies in Direct Coupling Analysis and related methods developed for multiple sequence alignments [74–76], which enabled structure determination for hundreds of protein families from metagenomic data [77] but require deep sequence alignments unavailable for orphan proteins and recently evolved sequences.

For protein language models, Zhang et al. developed the categorical Jacobian, which systematically mutates each residue to all 20 amino acids and measures changes in predicted probabilities across all positions [78]. Applied to ESM-2, this achieves 0.80 precision at L/2 for long-range contact prediction, surpassing mean-field DCA by 19% from single sequences. Critically, analysis of the learned dependencies revealed that ESM-2 stores coevolutionary statistics through “segment pair lookup” – matching local sequence motifs of approximately 22–30 residues rather than integrating information across full protein length. This finding is directly relevant to the mechanistic interpretability agenda: it suggests a specific computational strategy rather than abstract feature encoding.

For genomic models, Tomaz da Silva et al. introduced nucleotide dependency analysis across 14 models, achieving AUROC exceeding 0.9 for RNA secondary structure detection, 0.92 for pseudoknot identification, and 0.8 for tertiary contacts [8]. Dependency maps exhibit distinct signatures for different functional elements: diagonal blocks for regulatory motifs, off-diagonal blocks for interacting elements such as splice donor-acceptor pairs, and antiparallel diagonals for RNA secondary structure. The study demonstrated that multispecies training is essential for learning structural dependencies and experimentally validated four novel RNA structures in E. coli 5’UTR regions using DMS-MaPseq chemical probing.

Jacobian analysis is computationally expensive (O(L × A) forward passes) and, for genomic models, sensitive to tokenization artifacts [4]. Yet it occupies a unique position: where attention and attribution characterize individual inputs, Jacobian analysis extracts pairwise relationships from single sequences, bridging foundation models and classical coevolutionary analysis.

Table 1 summarizes the four approaches. Collectively, they establish that biological foundation models encode substantial structural and functional knowledge – residue contacts, secondary structure, regulatory elements, splice logic, and even RNA tertiary interactions are recoverable from representations learned through self-supervised training. Yet these methods share a fundamental explanatory limit [79]. Attention reveals what positions the model considers related, but these relationships can be altered without affecting predictions. Attribution quantifies which inputs drive outputs, but gradients in sequence space suffer from off-simplex artifacts and do not expose computational mechanisms. Probing confirms that biological properties are linearly decodable, but encoded information is not necessarily utilized during inference [9]. Even Jacobian analysis, which comes closest to mechanistic characterization through the “segment pair lookup” finding, operates on input-output relationships rather than internal representations.

**Table 1.** Comparison of post-hoc behavioral interpretation methods for biological foundation models.

| Method | Analytical Target | Output | Computational Scaling | Biological Validation Examples | Methodological Considerations |
| --- | --- | --- | --- | --- | --- |
| Attention Analysis | Pairwise token relationships learned by the model | Attention weight matrices [$L \times L$] per head per layer | $O(1)$ extraction; $O(L^2)$ storage per head | Residue-residue contacts in ESM [37]; splice site pairing in SpliceBERT [41] | Attention distributions can be altered without changing predictions [42]; information mixing across layers complicates single-layer interpretation [45] |
| Attribution Methods | Per-position contribution to a specific prediction | Importance scores per input position | $O(L)$ for ISM; $O(1)$ for gradient methods | Functionally important residues in Pab1 [53]; splice site motifs in GENA-LM [54] | Off-simplex gradients affect 10–20% of positions in sequence models [57]; some methods produce outputs independent of model parameters [55] |
| Probing Tasks | Information linearly decodable from representations | Classifier accuracy on frozen embeddings | $O(n)$ probe training; evaluation per layer | Secondary structure Q8=71.6% for ESM-1b [6]; splice site MCC=96% for Nucleotide Transformer [4] | Probe complexity can confound results [68]; high probe accuracy does not imply feature utilization by the model [9] |
| Jacobian Analysis | Pairwise dependencies between sequence positions | Dependency maps [$L \times L$] or tensors [$L \times A \times L \times A$] | $O(L \times A)$ forward passes | Contact prediction precision=0.80 for ESM-2 [78]; RNA structure AUROC>0.9 across 14 gLMs [8] | Requires categorical rather than numerical perturbations for neural models [78]; tokenization artifacts in k-mer models [4] |

The common thread is that behavioral methods characterize models from outside – they describe what goes in, what comes out, and what correlates with what, without revealing how information is organized and transformed within the network. Moving from behavioral characterization to understanding internal computational structure requires methods that decompose the representational space itself. Sparse dictionary learning provides such methods.

## 3. Sparse Dictionary Learning for Biological Foundation Models

The methods surveyed in Section 2 establish what biological foundation models encode but not how this information is internally organized – a gap rooted in the structure of neural network representations themselves. Individual neurons in both large language models and biological language models respond to multiple, semantically distinct concepts, a phenomenon termed polysemanticity. The superposition hypothesis proposes that this entanglement arises from a geometric property of high-dimensional spaces – a network with M neurons can represent N >> M distinct features by exploiting the fact that most features are sparse, rarely co-occurring in any given input [14]. This representational strategy is computationally efficient but renders individual neurons uninterpretable, as each participates in encoding multiple features simultaneously.

Sparse autoencoders (SAEs) address this by learning an overcomplete dictionary that decomposes dense, polysemantic activations into sparse, approximately monosemantic features. The approach trains an autoencoder whose hidden layer is expanded by a factor of 8–32× relative to the input dimension, with sparsity constraints ensuring that only a small subset of dictionary elements activates for any given input. If the superposition hypothesis is correct – if models indeed compress many sparse features into fewer dimensions – then an appropriately overcomplete, sparsity-constrained dictionary should recover the underlying feature directions.

An important distinction governs the interpretation of SAE results. Recovering features that correlate with biological annotations establishes representational interpretability – identifying which directions in activation space encode which concepts. This is distinct from computational interpretability – understanding how information is transformed between layers – and from causal mechanistic understanding – demonstrating that specific features or circuits are necessary and sufficient for model predictions. The vast majority of biological SAE work to date operates at the representational level. Transcoders approach the computational level by decomposing inter-layer transformations, and causal activation patching begins to address the causal level, but these remain isolated efforts rather than standard practice.

Notably, SAE-derived features and behavioral probing results are not independent. Early SAE studies on protein language models found that recovered features correspond to the same biological concepts detected by probing – secondary structures, functional domains, binding sites – but with substantially higher specificity. Where probing demonstrates that a representation contains information about α-helices, SAE decomposition identifies which directions in activation space encode helical structure and how those directions relate to other features. In this sense, SAE analysis provides a representational substrate for behavioral observations: the linear separability that probes detect can often be traced to specific SAE features [18,19]. Whether this correspondence is universal or reflects shared biases in evaluation methodology remains an open question (Section 4).

### 3.1 Sparse Autoencoder Methodology and Architectural Variants

All SAE architectures share a common structure: an encoder maps input activations x to a sparse hidden representation h, and a decoder reconstructs the original activations as 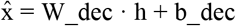. The optimization objective balances reconstruction fidelity (typically measured as 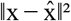) against sparsity of h. Architectures differ in how sparsity is enforced and what additional structure is imposed on the feature space. Table 2 provides formal definitions; Figure 2 illustrates the computational scheme. Below we describe the principal variants and their relevance to biological applications.

**Table 2.**
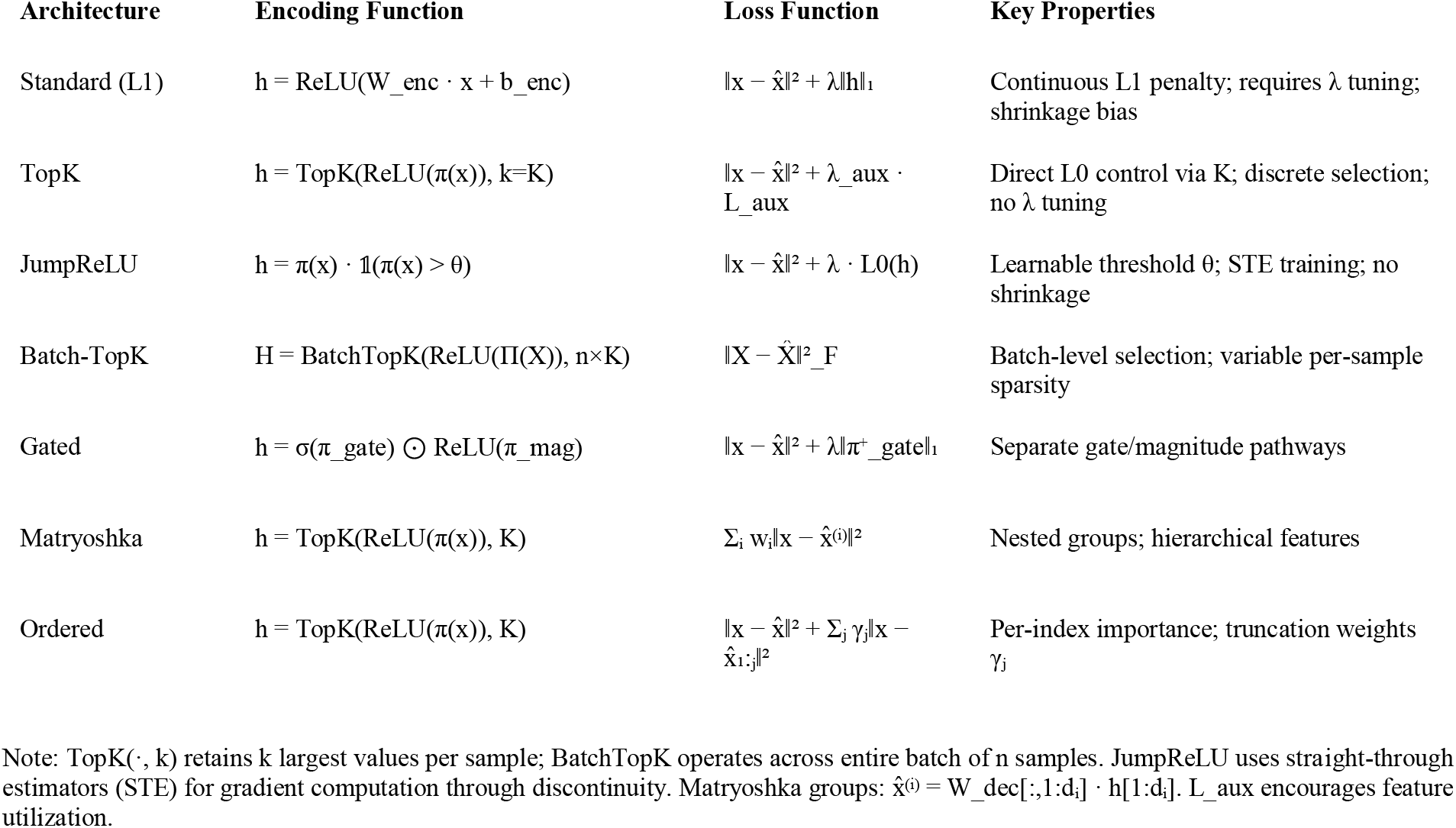
Sparse Autoencoder Architecture Comparison. All architectures share the form: encode input x → sparse representation h → reconstruction 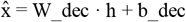. Notation: π(x) = W_enc · x + b_enc (pre-activation); σ (sigmoid); □ (indicator function); ⊙(element-wise product); ‖·‖_F (Frobenius norm).

**Figure 2.**
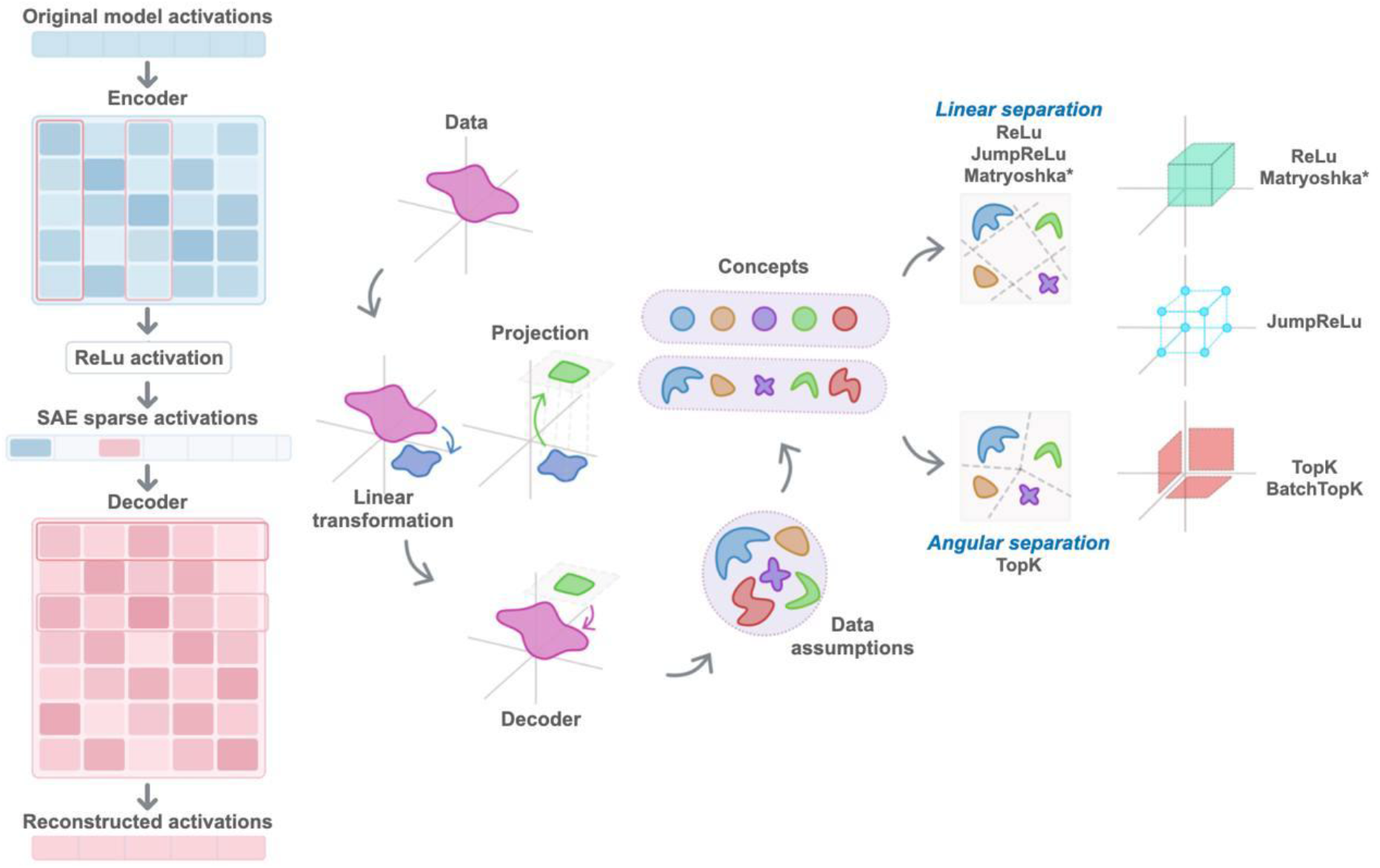
Sparse autoencoder architecture and the feature-to-experiment pipeline. *(Left)* Model activations (h ∈ ℝ^n^) are encoded into an overcomplete sparse representation (f ∈ ℝ^m^, m >> n) and reconstructed through a decoder; sparsity ensures only a small subset of features activates per input. *(Right)* Translational pipeline: SAE features extracted from a pretrained model generate biological hypotheses that can be tested through high-throughput assays (MPRA, CRISPR screens).

#### 3.1.1 From L1 Regularization to Direct Sparsity Control

The original SAE architecture enforces sparsity through an L1 penalty on the hidden activations: the loss function 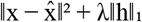 penalizes the sum of absolute activation values, encouraging most features to remain near zero [16] . ReLU activations ensure non-negativity, so each feature represents presence rather than absence of a pattern. This architecture was applied in early biological interpretability work, including pathology foundation models [25] and HyenaDNA genomic models [22]. However, the L1 penalty introduces two practical limitations: the sparsity coefficient λ requires careful tuning to achieve target sparsity levels, and the penalty induces shrinkage bias – uniformly attenuating all activations, including those of genuinely important features.

TopK SAEs replace the continuous L1 penalty with a discrete constraint: the TopK operation retains only the K largest activations per input and zeros all others [17]. This provides direct control over sparsity through K rather than requiring iterative tuning of λ, simplifies cross-model comparison at matched sparsity levels, and preserves the full magnitude of selected features, eliminating shrinkage bias. TopK has become the dominant architecture for protein language model interpretability, adopted for ESM-2 analysis across multiple studies [18– 20], as well as for fine-tuned protein models [80], antibody language models [81], and protein structure generation models [27].

JumpReLU SAEs take a different approach to sparsity control by introducing a learnable per-feature threshold θ: the activation function outputs π(x) · 𝟙 (π(x) > θ), where π(x) is the pre-activation and 𝟙 is the indicator function [82]. Training proceeds through straight-through estimators (STEs) that approximate gradients through the discontinuous threshold, enabling direct optimization of L0 sparsity – the count of active features – rather than relying on L1 as a proxy. Evaluation on Gemma 2 9B demonstrated that JumpReLU achieves state-of-the-art reconstruction fidelity at matched sparsity levels compared to both Gated and TopK architectures. JumpReLU is closely related to Gated SAEs [83], which separate feature selection from magnitude estimation through parallel sigmoid-gate and ReLU-magnitude pathways – a principled decomposition that permits independent representation of feature relevance and expression strength. With weight sharing the two architectures are mathematically equivalent, though they differ in training dynamics and loss formulation [82]. Neither Gated nor JumpReLU SAEs have been widely applied to biological models, though their adoption in Gemma Scope [84] as the default architecture for language model interpretability suggests they may see broader use as biological SAE analyses mature.

#### 3.1.2 Batch-Level and Hierarchical Architectures

The architectures above operate on individual samples. Batch-TopK SAEs modify the TopK operation to select the top n×K activations across an entire batch of n samples rather than independently per sample [85]. This allows variable per-sample sparsity: a sample containing a rare regulatory motif might activate more features, while a generic sample activates fewer. The batch-level selection proves particularly valuable when biologically important patterns appear infrequently in individual sequences but consistently across datasets – a common property of regulatory elements, resistance genes, and specialized protein domains. Batch-TopK SAEs have been particularly effective in genomic models, where features like transcription factor binding sites or viral regulatory elements may appear in a small fraction of sequences. Applications include Nucleotide Transformer analysis [23], Evo 2 genomic foundation model interpretation [24], and single-cell foundation models scGPT and scFoundation [26].

The architectures described so far impose no organization on the learned feature space. Matryoshka SAEs introduce hierarchical structure by organizing features into nested groups of increasing size (d_1_ < d_2_ < … < d_n), where each group must independently reconstruct the input using only its allocated feature subset [86]. The loss function incorporates weighted reconstruction terms for each group (Table 2). This progressive capacity constraint forces earlier groups to capture high-level, abstract patterns – because they have fewer features – while later groups encode finer details. Importantly, this hierarchy emerges from the training objective without explicit biological supervision; abstract features arise because they represent the most efficient way to minimize reconstruction error under severe capacity constraints. Application to ESM2-3B at layers 18 and 36 recovered features spanning coarse-grained domains and regions in early groups to fine-grained motifs and targeting peptides in later groups [87], and analysis of ESM-2 650M confirmed the potential for multi-scale feature organization [20].

Ordered SAEs extend hierarchical organization by assigning explicit per-index importance ordering with decreasing truncation weights, creating a continuous hierarchy rather than discrete groups [81]. Direct comparison with TopK SAEs in antibody language models revealed a fundamental trade-off: TopK SAEs produced sparse, localized features specific to complementarity-determining regions, facilitating discrete structural interpretation, while Ordered SAEs generated broader activation patterns enabling hierarchical steering through coarse-to-fine adjustments. Architecture selection thus depends on the application objective – interpretability of discrete biological elements versus multi-scale generative control.

### 3.2 Transcoders: Decomposing Computational Transformations

SAEs decompose representations at individual layers – they answer which concepts are encoded at a given computational stage. Transcoders address a complementary question: how models transform information between layers. Rather than reconstructing a single activation vector, a transcoder learns a sparse mapping from layer L input to layer L output (or from one layer to the next), decomposing the residual stream update into sparse latent features (Figure 2). This architecture, originally proposed for large language models [88], enables identification of computational circuits: specific pathways through which information propagates and is processed across network depth.

The distinction is substantive. An SAE at layer 10 of a protein language model might identify features corresponding to α-helices or functional domains. A transcoder analyzing the same computational block would reveal how the model constructs these representations – which lower-level features are composed to form higher-level ones, exposing the compositional structure of biological knowledge. Joint application of SAEs and transcoders to ESM-2 650M confirmed that the two approaches provide complementary insights: SAEs characterize what is represented, while transcoders reveal how representations are computed [20]. However, transcoder applications to biological models remain limited to this single study, and the biological interpretation of transcoder-derived circuits has not been systematically validated – an important gap given the method’s potential for revealing hierarchical biological computation.

### 3.3 Applications Across Biological Domains

Application of SAEs to biological foundation models has grown rapidly, spanning protein language models, genomic models, single-cell foundation models, pathology vision transformers, and specialized architectures for antibodies and protein structure generation. Table 3 provides a systematic overview of published studies. Rather than reviewing individual papers sequentially, we organize the discussion around cross-cutting patterns that emerge across domains.

**Table 3.**
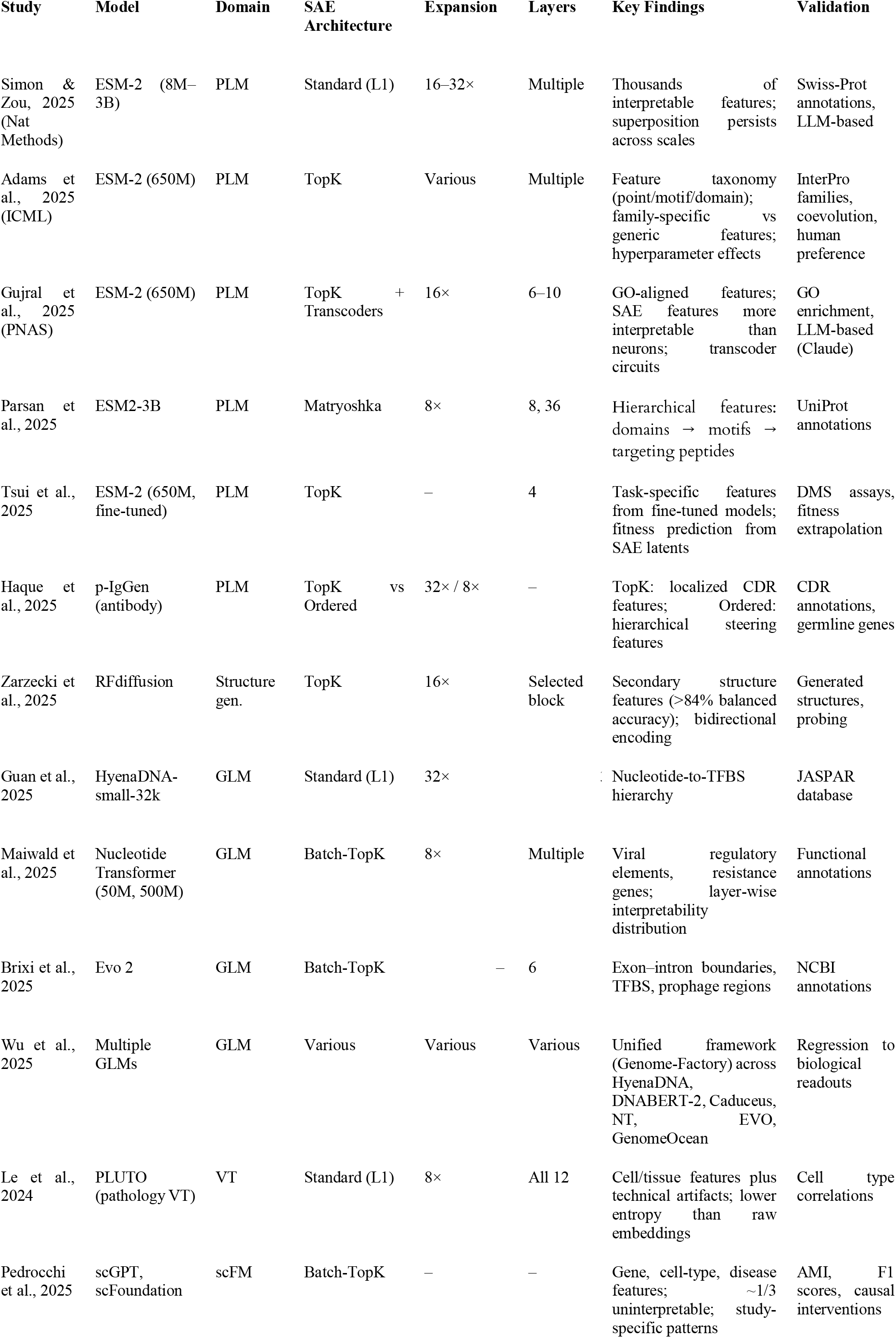
Overview of SAE studies applied to biological foundation models. PLM = protein language model; GLM = genomic language model; scFM = single-cell foundation model; VT = vision transformer.

#### 3.1.1 Feature Recovery and Biological Alignment

The most consistent finding across all SAE studies is that sparse decomposition recovers features aligned with established biological knowledge. In protein language models, multiple independent analyses of ESM-2 – using different SAE architectures, expansion factors, layers, and evaluation methods – converge on the same classes of biological features: secondary structures (α-helices, β-strands), conserved sequence motifs, functional domains, post-translational modification sites, localization signals, and biochemical properties [18–20]. This convergence across independent studies provides stronger evidence for the biological meaningfulness of SAE features than any single analysis could.

In genomic models, SAE features span a comparable hierarchy: from elementary patterns (individual nucleotide identity) through transcription factor binding sites to complex regulatory elements and multi-kilobase structural features. Standard SAEs applied to HyenaDNA recovered features ranging from single-nucleotide specificity to C2H2 zinc finger and MADS-box transcription factor binding sites, validated against the JASPAR database [22]. Batch-TopK SAEs trained on Nucleotide Transformer models identified viral regulatory elements (CMV enhancer, HIV-gag protein, Rev Response Element) and antibiotic resistance genes (kanamycin, streptomycin, puromycin resistance) in plasmid sequences [23]. Analysis of Evo 2 recovered features corresponding to exon–intron boundaries, transcription factor binding sites, and prophage regions, validated against NCBI genome annotations [24].

Beyond sequence models, SAE features capture domain-appropriate concepts. In pathology vision transformers (PLUTO), features spanned cellular morphology (carcinoma, red blood cells, mucin), tissue architecture (stromal clefts, fiber orientations), and – notably – technical artifacts (blur, sectioning artifacts, staining variations), demonstrating that SAEs decompose not only biological signal but also experimental confounds [25]. In single-cell foundation models, features organized into gene-specific patterns (immunoglobulins, metallothioneins, HLA genes), cell-type identities, and disease or technical variation (COVID-19 status, sequencing protocols), with approximately one-third of features difficult to interpret through manual annotation [26]. In antibody models, features corresponded to CDR identity and germline gene usage [81]. In the protein structure generation model RFdiffusion, individual features encoded secondary structure patterns with balanced accuracies exceeding 84%, and some features encoded both helix and strand identity through opposite activation directions [27].

Across all domains, SAE features are substantially more interpretable than individual neurons. Simon & Zou (2025) reported thousands of concept-aligned features per layer versus dozens for raw neurons [18]; Le et al. (2024) demonstrated lower entropy and stronger cell-type specificity for SAE features compared to raw embeddings [25]. This consistent improvement supports the superposition hypothesis: biological models compress many more features into their activation spaces than the number of available dimensions.

#### 3.3.2 Hierarchical Organization and Layer Specialization

Two forms of hierarchy emerge from SAE analyses: hierarchical organization within a layer (imposed by architecture) and hierarchical specialization across layers (emergent from model training).

Within-layer hierarchy is most explicitly captured by Matryoshka SAEs. Application to ESM2-3B revealed that early feature groups encode coarse-grained patterns – protein domains, broad structural regions – while later groups capture fine-grained motifs and targeting peptides [87]. This mirrors the multi-scale organization of protein biology and suggests that capacity-constrained feature learning naturally prioritizes structural hierarchy. Ordered SAEs provide a continuous variant, with high-importance indices capturing broad patterns and low-importance indices encoding specifics [81].

Across-layer specialization is reported consistently but with important methodological caveats. Adams et al. (2025) systematically varied layer choice and found that middle layers contain the most family-specific features [19]. Maiwald et al. (2025) observed non-uniform interpretability distribution in Nucleotide Transformer: few interpretable features in early layers, peaking in intermediate layers, then declining [23]. Simon & Zou (2025) found that larger ESM-2 models encode more interpretable features than smaller ones, and that feature complexity increases with depth [18]. Tsui et al. (2025) specifically targeted deep layers (layer 24) of fine-tuned ESM-2 models and recovered task-specific functional features, while earlier layers in the same models captured general structural patterns [80]. These observations suggest that feature interpretability and biological specificity vary substantially across model architectures and layers – but systematic comparisons across studies are complicated by differences in SAE architectures, expansion factors, evaluation methods, and interpretability criteria.

#### 3.3.3 Architecture–Domain Interactions

The choice of SAE architecture interacts with the data domain in ways that are beginning to be understood. TopK SAEs dominate protein language model studies, where features tend to be locally concentrated (a motif, a domain boundary, a binding site). Batch-TopK SAEs have emerged as preferred for genomic applications, where biologically important regulatory elements may appear in a small fraction of sequences but consistently across datasets – a frequency distribution that per-sample TopK selection might underrepresent. The same Batch-TopK preference appears in single-cell models, where gene expression is extremely sparse (most genes are silent in most cells) but co-expression patterns are consistent across cell populations.

Expansion factors exhibit a notable pattern: larger factors for smaller models (32× for ESM-2-8M) and smaller factors for larger models (8× for ESM2-3B), suggesting that dictionary size should scale with model capacity but not necessarily linearly. Pathology vision transformers employed conservative expansion (8×), possibly reflecting higher intrinsic dimensionality of visual compared to sequence representations. These trends are empirical and have not been systematically benchmarked – the relationship between expansion factor, model size, and feature quality remains undercharacterized.

#### 3.3.4 Practical Patterns and Methodological Convergence

Several practical patterns emerge across the literature. Feature taxonomy development has proceeded in parallel across groups: Adams et al. (2025) categorized features by activation pattern (point, motif, domain) and family specificity [19]; Gujral et al. (2025) developed a classification based on activation patterns (point, periodic, motif, domain features) [20]; both frameworks proved useful for systematic feature analysis. Validation approaches have similarly converged on a common repertoire: Gene Ontology enrichment analysis, annotation matching against curated databases (UniProt, JASPAR, InterPro, NCBI annotations), and automated interpretability using large language models. Genome-Factory [89] has formalized reproducible SAE methodology across genomic architectures, and InterProt [19] provides open-source visualization for feature exploration.

However, these converging methodologies also share limitations that should inform interpretation of the results above. Current validation approaches predominantly match SAE features against existing annotations – a strategy that confirms feature alignment with known biology but cannot assess whether features capture genuinely novel patterns or biologically meaningful distinctions not yet catalogued. Features annotated as “difficult to interpret” (approximately one-third in Pedrocchi et al., 2025 [26]) may represent technical artifacts, novel biology, or simply limitations of the annotation vocabulary. Cross-study reproducibility of specific features has not been systematically assessed: do independent SAE trainings on the same model recover the same features, or are the decompositions sensitive to training details? Furthermore, biological alignment based on annotation matching does not establish that SAE features capture the model’s actual computational strategy – they may represent statistically salient directions that correlate with but do not causally determine model behavior.

Sparse dictionary learning provides a principled framework for decomposing the internal representations of biological foundation models into interpretable components. The methodology has matured rapidly: from initial demonstrations that SAEs recover biologically meaningful features to systematic analyses spanning multiple model families, architectures, and data modalities. Architectural choices interact meaningfully with biological domains – TopK for protein models with locally concentrated features, Batch-TopK for genomic and single-cell models with rare but consistent patterns, Matryoshka and Ordered variants for applications requiring hierarchical organization or generative steering.

The consistency of biological feature recovery across independent studies, model sizes, and analysis frameworks provides strong evidence that biological foundation models encode interpretable structure that SAEs can access. At the same time, the almost exclusive reliance on annotation-matching validation, the unresolved question of feature reproducibility across SAE training runs, and the limited deployment of causal interventions to establish that recovered features drive model behavior rather than merely correlate with it – all point to significant gaps between current practice and the goal of mechanistic biological understanding. These limitations, and the research directions they motivate, are examined in detail in the next section.

## 4. Limitations and Future Directions

The preceding sections documented how sparse dictionary learning recovers biologically interpretable features from foundation models across multiple domains. Yet several open questions remain regarding the stability, interpretation, and validation of these features. Many originate in language model interpretability research but acquire particular significance when features are intended to inform biological understanding.

### 4.1 Feature Stability, Granularity, and the Limits of Decomposition

SAE training involves stochastic optimization, and the resulting feature dictionaries depend on random initialization. Paulo and Belrose quantified this dependence for SAEs trained on Llama 3 8B: at cosine similarity threshold 0.7, only approximately 30% of features were shared across runs differing solely in random seed [90]. TopK architectures exhibited greater seed dependence than L1-regularized variants. Whether similar variability affects biological model SAEs has not been systematically examined, though the underlying optimization dynamics are shared. This observation does not invalidate individual features but complicates claims about comprehensive feature catalogs – if 70% of features change between runs, the decomposition captures a sample from the space of valid sparse bases rather than a unique ground-truth dictionary.

Related work has examined whether SAE features constitute atomic units of representation. Chanin et al. described feature absorption: when hierarchically related concepts co-occur, sparsity pressure causes general features to fail activating in contexts where more specific features fire [91]. Leask et al. introduced meta-SAEs – autoencoders trained on SAE decoder matrices – and showed that features from larger SAEs decompose into sparse combinations of features from smaller SAEs: an “Einstein” feature, for instance, decomposed into meta-latents corresponding to “scientist,” “Germany,” and “famous person” [92]. These findings suggest that feature granularity depends on dictionary size and sparsity settings rather than reflecting intrinsic structure. For biological applications, the implication is that the appropriate level of decomposition – residues, motifs, domains, or functional modules – likely varies with analytical objectives, and a single SAE training run captures one projection of a continuous hierarchy rather than the hierarchy itself. Matryoshka and Ordered SAE architectures (Section 3.1.2) provide explicit hierarchical organization, though optimal configurations for specific biological questions remain to be determined.

A more fundamental challenge concerns what SAE features actually represent. Heap et al. demonstrated that SAEs trained on randomly initialized (untrained) transformers produce features with auto-interpretability scores comparable to those obtained from trained models [93]. This result does not imply that SAE features are meaningless – trained and random models differ in other evaluation dimensions – but it raises the possibility that a substantial fraction of recovered features reflect statistical regularities of the training data and architectural induction biases rather than learned computational strategies. For biological models trained on protein or genomic sequences, where the training data itself contains strong statistical structure (amino acid frequencies, codon usage, evolutionary conservation), distinguishing data-driven features from model-learned features is particularly important and has not been systematically addressed.

The reconstruction–sparsity tradeoff introduces a further limitation. SAEs trained on biological foundation models typically explain 90–95% of activation variance at moderate sparsity levels. The remaining residual is not random noise: Engels et al. demonstrated that approximately half of SAE reconstruction error can be linearly predicted from input activations, indicating systematic information loss [94]. This structured residual may encode features requiring more than K active dictionary elements, continuous properties poorly captured by discrete activations, or computational dimensions orthogonal to the learned basis. The practical consequence is that SAE analysis provides a partial view of model representations; features contributing to model behavior may reside outside the recovered dictionary.

Taken together, these findings reframe the status of SAE features. Rather than fixed atoms of representation, they are projections – influenced by dictionary size, sparsity constraints, random seed, and the inherent limitations of sparse linear approximation. This does not diminish their utility for hypothesis generation and biological discovery, but it places important constraints on how confidently specific features can be treated as definitive units of model computation.

### 4.2 The Validation Gap: From Annotation Correlation to Causal and Experimental Evidence

Evaluating SAE features requires distinguishing three levels of evidence, corresponding to the interpretability levels introduced in Section 3. Annotation matching establishes representational alignment – that feature directions correspond to known biological concepts. Causal interventions test computational relevance – that features participate in the model’s actual prediction mechanisms. Experimental validation grounds both levels in biology, confirming that computationally relevant features reflect genuine biological properties rather than dataset artifacts. Current biological SAE studies operate almost exclusively at the first level; the gap between representational alignment and mechanistic understanding structures the concerns below.

The annotation-matching paradigm establishes biological coherence (Section 3.3.4) but not computational relevance. Three specific failure modes deserve attention. First, annotation matching cannot distinguish whether the model explicitly represents an annotated concept or has learned correlated sequence patterns for unrelated computational reasons. Second, co-occurring annotations that share sequence signatures may be collapsed into a single feature, conflating distinct biological processes. Third, annotation databases are incomplete and systematically biased toward well-characterized systems in model organisms – features may appear uninterpretable because relevant annotations do not exist, or may achieve high alignment scores by detecting database artifacts rather than biological properties.

A further methodological gap concerns the absence of null models for biological SAE evaluation. Without baselines – SAEs trained on randomly initialized models, on models with shuffled training data, or evaluated against randomized annotations – absolute values of alignment metrics such as AMI or F1 cannot be meaningfully interpreted. The fraction of difficult-to-interpret features noted in Section 3.3.1 acquires different significance depending on what proportion a null model would also produce: if SAEs on random transformers yield a comparable proportion of ostensibly “interpretable” features, as Heap et al. (2025) [93] suggest is plausible for auto-interpretability scores, then annotation-matching metrics may substantially overestimate the biological content of learned representations. Establishing standard null models – analogous to shuffled-label controls in supervised learning – is a prerequisite for quantitative claims about SAE feature quality.

A related concern is circularity of validation. Protein language models are trained on sequences drawn from UniProt and related databases; SAE features extracted from these models are then validated against UniProt functional annotations, Gene Ontology terms, and InterPro domain assignments derived from the same sequence corpus. This creates a closed evaluation loop in which features may reflect memorized training set statistics – associations between sequence patterns and co-occurring annotations – rather than generalizable biological representations. The problem is not unique to SAE evaluation; it affects all benchmark-based assessment of biological foundation models, but acquires particular significance when SAE features are presented as evidence of learned biological knowledge. Circularity can be partially addressed by validation against experimental data not represented in training annotations – deep mutational scanning landscapes, MPRA measurements, or structural studies published after model training – but no published biological SAE study has explicitly controlled for this confound.

These concerns intersect with broader questions about what biological foundation models actually encode. Tang et al. evaluated genomic language model embeddings on regulatory prediction tasks and found that Nucleotide Transformer (2.5B parameters) achieved Pearson r = 0.125 on HepG2 MPRA data, compared to r = 0.510 for supervised Enformer and r = 0.324 for CNNs trained on one-hot encodings – performance the authors characterize as “only marginally more informative than bag-of-dinucleotide statistics” [71]. For protein models, Zhang et al. analyzed ESM-2 using categorical Jacobian methods and found that contact prediction relies on matching local sequence motifs of 22–30 residues rather than integrating information across full protein length – a strategy they term “segment pair lookup” [78]. If foundation models rely substantially on local statistical patterns, SAE features that correlate with biological annotations may reflect these patterns rather than deeper mechanistic representations. Importantly, these findings are not contradictory but operate at different explanatory levels. SAE features characterize representational geometry – which directions in activation space correlate with annotated biological concepts – whereas SPL describes a computational strategy for a specific prediction task. A model may represent domain-level patterns because they are statistically regular in training data while computing contact predictions through local motif matching. This distinction reinforces the separation between representational and computational interpretability introduced in Section 3: the existence of domain-scale SAE features does not entail domain-scale computation. Several interpretations accommodate both SAE findings and these observations: standard benchmarks may not probe what models learn most deeply; statistical pattern learning at sufficient scale may produce representations functionally equivalent to mechanistic understanding for specific tasks; or models encode mixtures of shallow and deep patterns with SAE analysis preferentially surfacing the latter. Distinguishing these possibilities requires task designs that separate statistical regularities from generalizable biological principles.

From the interpretability side, multiple systematic evaluations have tempered expectations for SAE utility on downstream tasks. The Google DeepMind mechanistic interpretability team reported that SAEs underperformed linear probes on out-of-distribution generalization, leading them to deprioritize fundamental SAE research [95]. AxBench, a large-scale benchmark for concept detection and model steering, found that prompting and fine-tuning outperform all representation-based methods including SAEs, which were not competitive on either task [96]. A reconciling perspective argues that these negative results are specific to acting on known concepts – detecting or steering predefined properties – whereas SAEs retain value for discovering unknown concepts, where no predefined target exists [97]. This distinction is directly relevant to biological applications, where the primary use case is exploratory discovery of what models have learned rather than manipulation toward predetermined biological concepts. Nevertheless, the consistent underperformance on tasks with ground truth should temper claims about the practical utility of SAE features beyond hypothesis generation.

Causal interventions – demonstrating that feature manipulation produces predicted changes in model output – provide stronger evidence for the functional relevance of SAE features. To date, such experiments remain rare in biological applications, but the available results are encouraging. Nainani et al. adapted causal activation patching to the protein language model setting, performing it in SAE latent space to identify the minimal circuit responsible for contact prediction in ESM-2 [98]. For two case study proteins, preserving only 0.015–0.022% of latent–token pairs was sufficient to maintain prediction accuracy above 70% of the clean baseline. The analysis revealed that early-layer motif detectors causally gate domain recognition in deeper layers – a specific, testable mechanistic claim. This represents the first circuits-style causal analysis for protein language models and demonstrates the feasibility of moving beyond correlation. Feature steering experiments provide complementary evidence: Parsan et al. (2025) manipulated features associated with solvent-accessible surface area in ESMFold, demonstrating causal control over structural properties; Garcia & Ansuini [21] steered ESM-2 toward generating zinc finger domain sequences; and Pedrocchi et al. (2025) showed that intervening on SAE features in single-cell foundation models could reduce batch effects while preserving biological signal [26]. These interventions demonstrate that at least some SAE features are causally linked to model behavior, though the fraction of biologically validated features relative to the total dictionary remains small.

Biological applications enjoy a validation advantage unavailable in language model interpretability: high-throughput experimental assays – deep mutational scanning, massively parallel reporter assays, CRISPR screens, ChIP-seq – provide ground truth fundamentally stronger than annotation matching and without analog in natural language processing. Connecting SAE features to these experimental modalities, including negative controls that test whether feature–annotation correlations survive experimental perturbation, represents one of the most promising and distinctive directions for biological SAE research. Concrete workflows for such validation are discussed in Section 4.3.

### 4.3 Scaling, Standardization, and Open Problems

Computational requirements constrain current applications. Published SAE analyses have reached ESM2-3B (Matryoshka SAEs at layers 18 and 36 [87]) and Evo 2 at 7B parameters (Batch-TopK at layer 26 [24]), but larger models – including ESM-3 (98B parameters) and multimodal architectures such as ESM-3’s combined sequence-structure representation – lack SAE analysis. Training costs scale with model size, dictionary width, and the number of layers analyzed; storing activations for large-scale training can require terabytes of storage. Architectural innovations developed for language models address these constraints: TopK SAEs with fused sparse-dense kernels achieve approximately 6× compute efficiency at scale [17]; Switch SAEs route activations to smaller expert dictionaries, reducing per-sample computation [99]; KronSAE employs Kronecker factorization to reduce encoder parameters by 40–55% under fixed compute budgets [100]. Adapting these methods to biological models – which differ from language models in sequence length distributions, vocabulary sizes, and attention patterns – requires implementation effort rather than methodological innovation.

Features learned from one model do not transfer to architecturally distinct models. Different training objectives, tokenization schemes, and architectural choices produce different internal representations, so a feature corresponding to zinc coordination in ESM-2 may have no direct analog in ESM-3, ProtTrans, or models trained with different masking strategies. Tokenization plays a particularly consequential role in this incompatibility. It determines the basic unit of representation and thus the activation space in which SAE features are defined. A k-mer tokenized genomic model encodes multiple nucleotides per embedding position; SAE features activating on specific k-mers are methodologically incommensurable with features from single-nucleotide models, where each position corresponds to one base. This is not merely a difference in learned representations but a structural incompatibility – the spaces have different dimensionality, different positional semantics, and different relationships between token identity and biological sequence – that precludes meaningful cross-architecture feature alignment without explicit mapping between tokenization schemes. This model specificity means that interpretability findings are local: understanding one model does not directly inform understanding of another, and separate SAE training and validation is required for each architecture of interest. Whether functionally equivalent features – representing the same biological concepts through different learned directions – emerge across independently trained models remains an open empirical question with implications for both interpretability methodology and our understanding of the biological representations these models learn.

Standardized benchmarks would enable systematic comparison across studies. SAEBench provides comprehensive evaluation for language model SAEs across reconstruction, sparsity, absorption, and downstream task metrics [101], but no equivalent exists for biological domains. A biological SAEBench would need to incorporate domain-specific evaluation criteria: alignment with curated biological annotations, recovery of known functional sites, prediction of mutational effects, and – crucially – experimental validation benchmarks that go beyond computational metrics. The Genome-Factory framework [89] represents a step toward reproducible methodology across genomic architectures, but standardized evaluation across protein, genomic, and single-cell domains remains undeveloped.

Finally, supervised concept-based methods offer complementary approaches to SAE-driven discovery. Concept Activation Vectors (CAVs) identify directions in activation space corresponding to predefined concepts by training linear classifiers on concept-positive and concept-negative examples, with TCAV extending this to quantify concept influence on predictions [102]. CAVs require no architectural modification, making them applicable to any pretrained biological model, but test only for concepts specified in advance. Concept Bottleneck Protein Language Models go further by constraining neurons to represent predefined biological concepts, guaranteeing coverage of specified concepts at the cost of open-ended discovery [103]. For biological foundation models, where well-characterized properties coexist with unknown learned representations, these supervised methods and SAEs address complementary questions: whether a specific concept is encoded versus what concepts the model has learned. SAEs serve exploratory analysis; CAVs and concept bottlenecks serve applications requiring guaranteed alignment with specific biological ontologies.

Despite these open methodological questions, SAE-based interpretability already enables several translational applications. The most immediately actionable is model auditing: SAE features corresponding to technical artifacts – blur and sectioning artifacts in pathology models [25], batch effects and sequencing protocol signatures in single-cell models [26] provide concrete targets for detecting and removing unwanted dependencies before deployment. For regulatory applications – clinical variant interpretation, drug safety assessment – the ability to identify which biological properties drive a model’s predictions may prove as valuable as the predictions themselves, providing mechanistic transparency that black-box models cannot offer. The steering interventions described in Section 4.2 – structural control through surface area features [87], zinc finger domain generation [21], CDR diversification in antibody models [81] – demonstrate that SAE features can serve as interpretable control knobs for sequence design. However, no published steering experiment has yet been validated by expressing and functionally characterizing the designed sequences, leaving the gap between in silico feature manipulation and experimental outcomes unquantified.

A further promising direction concerns antimicrobial resistance surveillance. SAE features in Nucleotide Transformer models identify antibiotic resistance genes in plasmid sequences as discrete, interpretable features [23], and prophage region features recovered from Evo 2 [24] suggest analogous applications in phage genomics. If these features generalize beyond their training contexts, SAE-based decomposition could complement existing resistance gene databases by detecting novel or chimeric determinants that annotation-based methods miss – a capability of particular value as horizontal gene transfer continues to generate resistance mechanisms absent from curated catalogs.

The highest-impact but most distant application is hypothesis generation for experimental biology. The approximately one-third of SAE features that resist annotation matching [25,26] may include genuinely novel functional categories alongside decomposition artifacts. Distinguishing these requires connecting features to experimental assays through explicit workflows: for protein models, identifying features that activate consistently on unannotated positions across protein families and testing predicted functional residues through deep mutational scanning; for genomic models, nominating putative regulatory elements for validation via massively parallel reporter assays or CRISPR interference; for single-cell models, testing feature-derived co-expression modules through perturbation experiments. Tsui et al. (2025) provide a partial precedent by correlating SAE latent activations with DMS fitness landscapes [80], though the study validated known functional sites rather than discovering new ones.

Across these applications, a consistent pattern emerges: translational readiness decreases with the level of interpretability required. Model auditing operates at the representational level – identifying which features correspond to known artifacts – and is closest to routine deployment. Feature steering requires reproducible feature–output relationships and has reached proof-of-concept stage. Hypothesis generation demands causal mechanistic understanding validated experimentally and remains largely prospective. Biological domains hold a distinctive advantage in traversing this gradient: the availability of high-throughput experimental assays – DMS, MPRA, Perturb-seq, CRISPR screens – provides ground truth fundamentally stronger than annotation matching and without analog in natural language processing. Developing systematic frameworks that connect SAE feature discovery to these experimental validation pipelines represents the most promising path from interpretability as an analytical exercise to interpretability as a tool for biological discovery.

## 5. Conclusion

Sparse dictionary learning has shifted the interpretability of biological foundation models from behavioral characterization – establishing what information models encode – toward representational decomposition – identifying how that information is organized in activation space. The methods surveyed in this review document a rapid and consistent finding: SAEs recover features aligned with biological knowledge across protein language models, genomic models, single-cell foundation models, and pathology vision transformers, with convergence across independent studies providing stronger evidence than any single analysis. At the same time, this review has argued that a significant distance separates representational feature recovery from mechanistic understanding, and that the field’s reliance on annotation matching as a primary validation strategy leaves fundamental questions unresolved.

The three-level framework introduced in Section 3 clarifies the current state. Representational interpretability – identifying which directions in activation space correspond to biological concepts – is now supported by dozens of studies spanning multiple model families and data modalities. Computational interpretability – understanding how information is transformed between layers – has been addressed in a single study through transcoders applied to ESM-2. Causal mechanistic understanding – demonstrating that specific features or circuits are necessary and sufficient for predictions – has one proof of concept through activation patching in SAE latent space. The fraction of biologically validated features relative to the total dictionaries produced across all published studies remains small. This distribution is not a failure but a natural trajectory for a field less than two years old; it does, however, define where effort is most needed.

Four priorities emerge from the limitations identified in this review. First, null models and experimental validation should become standard practice: SAE features validated only against the same annotation databases used to train the underlying models cannot support claims of learned biological knowledge, and the absence of baselines renders alignment metrics uninterpretable. Second, transcoders and causal activation patching represent the methodological frontier for advancing beyond representational cataloging toward computational and causal understanding. Third, scaling SAE analysis to larger and multimodal architectures – ESM-3, Evo 2 at full scale, combined sequence-structure models – will test whether the patterns observed in current studies generalize or are artifacts of model size and architecture. Fourth, a standardized biological SAEBench incorporating domain-specific evaluation criteria – annotation alignment, functional site recovery, mutational effect prediction, and experimental benchmarks – would enable the systematic cross-study comparison that the field currently lacks.

Biological SAE interpretability sits at an inflection point. Sufficient evidence exists to establish that foundation models encode interpretable structure accessible through sparse decomposition; insufficient evidence exists to claim that this structure reveals how models compute biological predictions. What distinguishes biological applications from their NLP counterparts – and what may ultimately determine whether SAE analysis becomes a tool for genuine biological discovery – is the availability of experimental validation through deep mutational scanning, massively parallel reporter assays, and structural characterization. These techniques provide ground truth unavailable in language model interpretability. Whether the field invests in connecting SAE features to experimental evidence, rather than accumulating further annotation-matching results, will determine whether sparse dictionary learning advances biological understanding or remains a descriptive exercise.

## Acknowledgements

This work was supported by the Russian Science Foundation (grant No. 22-74-10053-P).

## Notes

### Competing Interest Statement

The authors have declared no competing interest.

## References

1. Simon, E.; Swanson, K.; Zou, J. Language Models for Biological Research: A Primer. Nat Methods 2024, 21, 1429–, doi:10.1038/s41592-024-02354-y.

2. Moor, M.; Banerjee, O.; Abad, Z.S.H.; Krumholz, H.M.; Leskovec, J.; Topol, E.J.; Rajpurkar, P. Foundation Models for Generalist Medical Artificial Intelligence. Nature 2023, 616, 265–, doi:10.1038/s41586-023-05881-4.

3. Lin, Z.; Akin, H.; Rao, R.; Hie, B.; Zhu, Z.; Lu, W.; Smetanin, N.; Verkuil, R.; Kabeli, O.; Shmueli, Y.; et al. Evolutionary-Scale Prediction of Atomic-Level Protein Structure with a Language Model. Science 2023, 379, 1130–, doi:10.1126/science.ade2574.

4. Dalla-Torre, H.; Gonzalez, L.; Mendoza-Revilla, J.; Lopez Carranza, N.; Grzywaczewski, A.H.; Oteri, F.; Dallago, C.; Trop, E.; de Almeida, B.P.; Sirelkhatim, H.; et al. Nucleotide Transformer: Building and Evaluating Robust Foundation Models for Human Genomics. Nat Methods 2025, 22, 297–, doi:10.1038/s41592-024-02523-z.

5. Elnaggar, A.; Heinzinger, M.; Dallago, C.; Rehawi, G.; Wang, Y.; Jones, L.; Gibbs, T.; Feher, T.; Angerer, C.; Steinegger, M.; et al. ProtTrans: Toward Understanding the Language of Life Through Self-Supervised Learning. IEEE Transactions on Pattern Analysis and Machine Intelligence 2022, 44, 7127–, doi:10.1109/TPAMI.2021.3095381.

6. Rives, A.; Meier, J.; Sercu, T.; Goyal, S.; Lin, Z.; Liu, J.; Guo, D.; Ott, M.; Zitnick, C.L.; Ma, J.; et al. Biological Structure and Function Emerge from Scaling Unsupervised Learning to 250 Million Protein Sequences. Proc Natl Acad Sci U S A 2021, 118, e2016239118, doi:10.1073/pnas.2016239118.

7. Avsec, Ž.; Agarwal, V.; Visentin, D.; Ledsam, J.R.; Grabska-Barwinska, A.; Taylor, K.R.; Assael, Y.; Jumper, J.; Kohli, P.; Kelley, D.R. Effective Gene Expression Prediction from Sequence by Integrating Long-Range Interactions. Nat Methods 2021, 18, 1203–, doi:10.1038/s41592-021-01252-x.

8. Tomaz da Silva, P.; Karollus, A.; Hingerl, J.; Galindez, G.S.T.; Wagner, N.; Hernandez-Alias, X.; Incarnato, D.; Gagneur, J. Nucleotide Dependency Analysis of Genomic Language Models Detects Functional Elements. Nat Genet 2025, 57, 2602–, doi:10.1038/s41588-025-02347-3.

9. Belinkov, Y. Probing Classifiers: Promises, Shortcomings, and Advances. Computational Linguistics 2022, 48, 219–, doi:10.1162/coli_a_00422.

10. Vig, J.; Madani, A.; Varshney, L.R.; Xiong, C.; Socher, R.; Rajani, N.F. BERTology Meets Biology: Interpreting Attention in Protein Language Models 2021.

11. Sundararajan, M.; Taly, A.; Yan, Q. Axiomatic Attribution for Deep Networks. In Proceedings of the Proceedings of the 34th International Conference on Machine Learning; PMLR, July 17 2017; pp. 3319–3328.

12. Biswas, S.; Khimulya, G.; Alley, E.C.; Esvelt, K.M.; Church, G.M. Low-N Protein Engineering with Data-Efficient Deep Learning. Nat Methods 2021, 18, 396–, doi:10.1038/s41592-021-01100-y.

13. Elhage, N.; Nanda, N.; Olsson, C.; Henighan, T.; Joseph, N.; Mann, B.; Askell, A.; Bai, Y.; Chen, A.; Conerly, T.; et al. A Mathematical Framework for Transformer Circuits; 2021.

14. Elhage, N.; Hume, T.; Olsson, C.; Schiefer, N.; Henighan, T.; Kravec, S.; Hatfield-Dodds, Z.; Lasenby, R.; Drain, D.; Chen, C.; et al. Toy Models of Superposition 2022.

15. Cunningham, H.; Ewart, A.; Riggs, L.; Huben, R.; Sharkey, L.Sparse Autoencoders Find Highly Interpretable Features in Language Models 2023.

16. Bricken, T.; Templeton, A.; Batson, J.; Chen, B.; Jermyn, A.; Conerly, T.; Turner, N.; Anil, C.; Denison, C.; Askell, A. Towards Monosemanticity: Decomposing Language Models with Dictionary Learning. Transformer Circuits Thread 2023, 2, 6.

17. Gao, L.; Tour, T.D. la; Tillman, H.; Goh, G.; Troll, R.; Radford, A.; Sutskever, I.; Leike, J.; Wu, J. Scaling and Evaluating Sparse Autoencoders 2024.

18. Simon, E.; Zou, J. InterPLM: Discovering Interpretable Features in Protein Language Models via Sparse Autoencoders. Nature methods 2025, 22, 2117–.

19. Adams, E.; Bai, L.; Lee, M.; Yu, Y.; AlQuraishi, M. From Mechanistic Interpretability to Mechanistic Biology: Training, Evaluating, and Interpreting Sparse Autoencoders on Protein Language Models 2025, 2025.02.06.636901.

20. Gujral, O.; Bafna, M.; Alm, E.; Berger, B. Sparse Autoencoders Uncover Biologically Interpretable Features in Protein Language Model Representations. Proc. Natl. Acad. Sci. U.S.A. 2025, 122, e2506316122, doi:10.1073/pnas.2506316122.

21. Garcia, E.N.V.; Ansuini, A. Interpreting and Steering Protein Language Models through Sparse Autoencoders 2025.

22. Guan, H.; He, J.; Zhang, J. Sparse Autoencoders Reveal Interpretable Structure in Small Gene Language Models 2025.

23. Maiwald, A.; Jedryszek, P.; Draye, F.; Schölkopf, B.; Morris, G.M.; Crook, O.M. Decode-Glm: Tools to Interpret, Audit, and Steer Genomic Language Models. bioRxiv 2025, 2025–10.

24. Brixi, G.; Durrant, M.G.; Ku, J.; Poli, M.; Brockman, G.; Chang, D.; Gonzalez, G.A.; King, S.H.; Li, D.B.; Merchant, A.T. Genome Modeling and Design across All Domains of Life with Evo 2. BioRxiv 2025, 2025–02.

25. Le, N.M.; Shen, C.; Patel, N.; Shah, C.; Sanghavi, D.; Martin, B.; Eng, A.; Shenker, D.; Padigela, H.; Biju, R.; et al. Learning Biologically Relevant Features in a Pathology Foundation Model Using Sparse Autoencoders 2024.

26. Pedrocchi, F.; Barkmann, F.; Joudaki, A.; Boeva, V. Sparse Autoencoders Reveal Interpretable Features in Single-Cell Foundation Models. bioRxiv 2025, 2025–10.

27. Zarzecki, W.; Szymczak, P.; Szczurek, E.; Deja, K. FoldSAE: Learning to Steer Protein Folding Through Sparse Representations 2025.

28. Lopez, R.; Regier, J.; Cole, M.B.; Jordan, M.I.; Yosef, N. Deep Generative Modeling for Single-Cell Transcriptomics. Nat Methods 2018, 15, 1058–, doi:10.1038/s41592-018-0229-2.

29. Li, Q.; Hu, Z.; Wang, Y.; Li, L.; Fan, Y.; King, I.; Jia, G.; Wang, S.; Song, L.; Li, Y. Progress and Opportunities of Foundation Models in Bioinformatics. Briefings in Bioinformatics 2024, 25, bbae548.

30. Guo, F.; Guan, R.; Li, Y.; Liu, Q.; Wang, X.; Yang, C.; Wang, J. Foundation Models in Bioinformatics. National science review 2025, 12, nwaf028.

31. Liu, X.; Zhang, Y.; Lu, Y.; Yin, C.; Hu, X.; Liu, X.; Chen, L.; Wang, S.; Rodriguez, A.; Yao, H. Biomedical Foundation Model: A Survey. arXiv preprint arXiv:2503.02104 2025.

32. Shu, D.; Wu, X.; Zhao, H.; Rai, D.; Yao, Z.; Liu, N.; Du, M. A Survey on Sparse Autoencoders: Interpreting the Internal Mechanisms of Large Language Models. arXiv preprint arXiv:2503.05613 2025.

33. Grešová, K.; Vaculík, O.; Alexiou, P. Using Attribution Sequence Alignment to Interpret Deep Learning Models for miRNA Binding Site Prediction. Biology 2023, 12, doi:10.3390/biology12030369.

34. Vig, J. A Multiscale Visualization of Attention in the Transformer Model. In Proceedings of the Proceedings of the 57th Annual Meeting of the Association for Computational Linguistics: System Demonstrations; Association for Computational Linguistics: Florence, Italy, 2019; pp. 37–42.

35. Goldowsky-Dill, N.; Chughtai, B.; Heimersheim, S.; Hobbhahn, M. Detecting Strategic Deception Using Linear Probes 2025.

36. Shi, Z.; Meng, Z.; Tuo, H.; Tan, C. Attribution-Based Interpretable Classification Neural Network with Global and Local Perspectives. Sci Rep 2025, 15, 24716, doi:10.1038/s41598-025-06218-z.

37. Rao, R.; Meier, J.; Sercu, T.; Ovchinnikov, S.; Rives, A. Transformer Protein Language Models Are Unsupervised Structure Learners 2020.

38. Rao, R.; Liu, J.; Verkuil, R.; Meier, J.; Canny, J.F.; Abbeel, P.; Sercu, T.; Rives, A. MSA Transformer 2021.

39. Lupo, U.; Sgarbossa, D.; Bitbol, A.-F. Protein Language Models Trained on Multiple Sequence Alignments Learn Phylogenetic Relationships. Nat Commun 2022, 13, 6298, doi:10.1038/s41467-022-34032-y.

40. Ji, Y.; Zhou, Z.; Liu, H.; Davuluri, R.V. DNABERT: Pre-Trained Bidirectional Encoder Representations from Transformers Model for DNA-Language in Genome. Bioinformatics 2021, 37, 2120–, doi:10.1093/bioinformatics/btab083.

41. Chen, K.; Zhou, Y.; Ding, M.; Wang, Y.; Ren, Z.; Yang, Y. Self-Supervised Learning on Millions of Primary RNA Sequences from 72 Vertebrates Improves Sequence-Based RNA Splicing Prediction. Briefings in Bioinformatics 2024, 25, bbae163, doi:10.1093/bib/bbae163.

42. Jain, S.; Wallace, B.C. [No Title Found]. In Proceedings of the Proceedings of the 2019 Conference of the North; Association for Computational Linguistics: Minneapolis, Minnesota, 2019; pp. 3543–3556.

43. Wiegreffe, S.; Pinter, Y. Attention Is Not Not Explanation. In Proceedings of the Proceedings of the 2019 Conference on Empirical Methods in Natural Language Processing and the 9th International Joint Conference on Natural Language Processing (EMNLP-IJCNLP); Association for Computational Linguistics: Hong Kong, China, 2019; pp. 11–20.

44. Bibal, A.; Cardon, R.; Alfter, D.; Wilkens, R.; Wang, X.; François, T.; Watrin, P. Is Attention Explanation? An Introduction to the Debate. In Proceedings of the Proceedings of the 60th Annual Meeting of the Association for Computational Linguistics (Volume 1: Long Papers); Association for Computational Linguistics: Dublin, Ireland, 2022; pp. 3889–3900.

45. Abnar, S.; Zuidema, W. Quantifying Attention Flow in Transformers. In Proceedings of the Proceedings of the 58th Annual Meeting of the Association for Computational Linguistics; Association for Computational Linguistics: Online, 2020; pp. 4190–4197.

46. Kobayashi, G.; Kuribayashi, T.; Yokoi, S.; Inui, K. Attention Is Not Only a Weight: Analyzing Transformers with Vector Norms. In Proceedings of the Proceedings of the 2020 Conference on Empirical Methods in Natural Language Processing (EMNLP); Association for Computational Linguistics: Online, 2020; pp. 7057–7075.

47. Lundberg, S.; Lee, S.-I. A Unified Approach to Interpreting Model Predictions 2017.

48. Shrikumar, A.; Greenside, P.; Kundaje, A. Learning Important Features Through Propagating Activation Differences 2019.

49. Zhou, J.; Troyanskaya, O.G. Predicting Effects of Noncoding Variants with Deep Learning–Based Sequence Model. Nat Methods 2015, 12, 934–, doi:10.1038/nmeth.3547.

50. Novakovsky, G.; Dexter, N.; Libbrecht, M.W.; Wasserman, W.W.; Mostafavi, S. Obtaining Genetics Insights from Deep Learning via Explainable Artificial Intelligence. Nat Rev Genet 2023, 24, 137–, doi:10.1038/s41576-022-00532-2.

51. Eraslan, G.; Avsec, Ž.; Gagneur, J.; Theis, F.J. Deep Learning: New Computational Modelling Techniques for Genomics. Nat Rev Genet 2019, 20, 403–, doi:10.1038/s41576-019-0122-6.

52. Meier, J.; Rao, R.; Verkuil, R.; Liu, J.; Sercu, T.; Rives, A. Language Models Enable Zero-Shot Prediction of the Effects of Mutations on Protein Function. Advances in neural information processing systems 2021, 34, 29303–.

53. Gelman, S.; Fahlberg, S.A.; Heinzelman, P.; Romero, P.A.; Gitter, A. Neural Networks to Learn Protein Sequence–Function Relationships from Deep Mutational Scanning Data. Proc. Natl. Acad. Sci. U.S.A. 2021, 118, e2104878118, doi:10.1073/pnas.2104878118.

54. Fishman, V.; Kuratov, Y.; Shmelev, A.; Petrov, M.; Penzar, D.; Shepelin, D.; Chekanov, N.; Kardymon, O.; Burtsev, M. GENA-LM: A Family of Open-Source Foundational DNA Language Models for Long Sequences. Nucleic Acids Research 2025, 53, gkae1310, doi:10.1093/nar/gkae1310.

55. Adebayo, J.; Gilmer, J.; Muelly, M.; Goodfellow, I.; Hardt, M.; Kim, B. Sanity Checks for Saliency Maps 2020.

56. Ghorbani, A.; Abid, A.; Zou, J. Interpretation of Neural Networks Is Fragile. In Proceedings of the Proceedings of the AAAI conference on artificial intelligence; 2019; Vol. 33, pp. 3681–3688.

57. Majdandzic, A.; Rajesh, C.; Koo, P.K. Correcting Gradient-Based Interpretations of Deep Neural Networks for Genomics. Genome Biol 2023, 24, 109, doi:10.1186/s13059-023-02956-3.

58. Prakash, E.I.; Shrikumar, A.; Kundaje, A. Towards More Realistic Simulated Datasets for Benchmarking Deep Learning Models in Regulatory Genomics. In Proceedings of the Machine Learning in Computational Biology; PMLR, 2022; pp. 58–77.

59. Jha, A.; K. Aicher, J.; R. Gazzara, M.; Singh, D.; Barash, Y. Enhanced Integrated Gradients: Improving Interpretability of Deep Learning Models Using Splicing Codes as a Case Study. Genome Biol 2020, 21, 149, doi:10.1186/s13059-020-02055-7.

60. Nair, S.; Shrikumar, A.; Schreiber, J.; Kundaje, A. fastISM: Performant in Silico Saturation Mutagenesis for Convolutional Neural Networks. Bioinformatics 2022, 38, 2403–, doi:10.1093/bioinformatics/btac135.

61. Schreiber, J.; Nair, S.; Balsubramani, A.; Kundaje, A. Accelerating in Silico Saturation Mutagenesis Using Compressed Sensing. Bioinformatics 2022, 38, 3564–, doi:10.1093/bioinformatics/btac385.

62. Conneau, A.; Kruszewski, G.; Lample, G.; Barrault, L.; Baroni, M. What You Can Cram into a Single $&!#* Vector: Probing Sentence Embeddings for Linguistic Properties. In Proceedings of the Proceedings of the 56th Annual Meeting of the Association for Computational Linguistics (Volume 1: Long Papers); Association for Computational Linguistics: Melbourne, Australia, 2018; pp. 2126–2136.

63. Belinkov, Y.; Durrani, N.; Dalvi, F.; Sajjad, H.; Glass, J. What Do Neural Machine Translation Models Learn about Morphology? In Proceedings of the Proceedings of the 55th Annual Meeting of the Association for Computational Linguistics (Volume 1: Long Papers); Association for Computational Linguistics: Vancouver, Canada, 2017; pp. 861–872.

64. Rao, R.; Bhattacharya, N.; Thomas, N.; Duan, Y.; Chen, P.; Canny, J.; Abbeel, P.; Song, Y. Evaluating Protein Transfer Learning with TAPE. Advances in neural information processing systems 2019, 32.

65. Zhou, Z.; Ji, Y.; Li, W.; Dutta, P.; Davuluri, R.; Liu, H. DNABERT-2: Efficient Foundation Model and Benchmark For Multi-Species Genome 2024.

66. Marin, F.I.; Teufel, F.; Horlacher, M.; Madsen, D.; Pultz, D.; Winther, O.; Boomsma, W. BEND: Benchmarking DNA Language Models on Biologically Meaningful Tasks 2024.

67. Nguyen, E.; Poli, M.; Faizi, M.; Thomas, A.; Birch-Sykes, C.; Wornow, M.; Patel, A.; Rabideau, C.; Massaroli, S.; Bengio, Y.; et al. HyenaDNA: Long-Range Genomic Sequence Modeling at Single Nucleotide Resolution 2023.

68. Hewitt, J.; Liang, P. Designing and Interpreting Probes with Control Tasks. In Proceedings of the Proceedings of the 2019 Conference on Empirical Methods in Natural Language Processing and the 9th International Joint Conference on Natural Language Processing (EMNLP-IJCNLP); Association for Computational Linguistics: Hong Kong, China, 2019; pp. 2733–2743.

69. Voita, E.; Titov, I. Information-Theoretic Probing with Minimum Description Length. In Proceedings of the Proceedings of the 2020 Conference on Empirical Methods in Natural Language Processing (EMNLP); Association for Computational Linguistics: Online, 2020; pp. 183–196.

70. Elazar, Y.; Ravfogel, S.; Jacovi, A.; Goldberg, Y. Amnesic Probing: Behavioral Explanation with Amnesic Counterfactuals. Transactions of the Association for Computational Linguistics 2021, 9, 175–, doi:10.1162/tacl_a_00359.

71. Tang, Z.; Somia, N.; Yu, Y.; Koo, P.K. Evaluating the Representational Power of Pre-Trained DNA Language Models for Regulatory Genomics 2024.

72. Sarkar, A.; Kang, Y.; Somia, N.; Puccetti, P.M.; Zhou, J.; Nagai, M.; Tang, Z.; Zhao, C.; Koo, P.K. Designing DNA With Tunable Regulatory Activity Using Score-Entropy Discrete Diffusion 2024.

73. Csendes, G.; Sanz, G.; Szalay, K.Z.; Szalai, B. Benchmarking Foundation Cell Models for Post-Perturbation RNA-Seq Prediction. BMC Genomics 2025, 26, 393, doi:10.1186/s12864-025-11600-2.

74. Kamisetty, H.; Ovchinnikov, S.; Baker, D. Assessing the Utility of Coevolution-Based Residue–Residue Contact Predictions in a Sequence- and Structure-Rich Era. Proc. Natl. Acad. Sci. U.S.A. 2013, 110, 15679–, doi:10.1073/pnas.1314045110.

75. Morcos, F.; Pagnani, A.; Lunt, B.; Bertolino, A.; Marks, D.S.; Sander, C.; Zecchina, R.; Onuchic, J.N.; Hwa, T.; Weigt, M. Direct-Coupling Analysis of Residue Coevolution Captures Native Contacts across Many Protein Families. Proc. Natl. Acad. Sci. U.S.A. 2011, 108, doi:10.1073/pnas.1111471108.

76. Ekeberg, M.; Lövkvist, C.; Lan, Y.; Weigt, M.; Aurell, E. Improved Contact Prediction in Proteins: Using Pseudolikelihoods to Infer Potts Models. Phys. Rev. E 2013, 87, 012707, doi:10.1103/PhysRevE.87.012707.

77. Ovchinnikov, S.; Park, H.; Varghese, N.; Huang, P.-S.; Pavlopoulos, G.A.; Kim, D.E.; Kamisetty, H.; Kyrpides, N.C.; Baker, D. Protein Structure Determination Using Metagenome Sequence Data. Science 2017, 355, 298–, doi:10.1126/science.aah4043.

78. Zhang, Z.; Wayment-Steele, H.K.; Brixi, G.; Wang, H.; Kern, D.; Ovchinnikov, S. Protein Language Models Learn Evolutionary Statistics of Interacting Sequence Motifs. Proc. Natl. Acad. Sci. U.S.A. 2024, 121, e2406285121, doi:10.1073/pnas.2406285121.

79. Rai, D.; Zhou, Y.; Feng, S.; Saparov, A.; Yao, Z. A Practical Review of Mechanistic Interpretability for Transformer-Based Language Models 2025.

80. Tsui, D.; Talreja, K.; Aghazadeh, A. Sparse Autoencoders for Low-$N$ Protein Function Prediction and Design 2025.

81. Haque, R.; Turnbull, O.M.; Parsan, A.; Parsan, N.; Yang, J.J.; Deane, C.M. Mechanistic Interpretability of Antibody Language Models Using SAEs 2025.

82. Rajamanoharan, S.; Lieberum, T.; Sonnerat, N.; Conmy, A.; Varma, V.; Kramár, J.; Nanda, N. Jumping Ahead: Improving Reconstruction Fidelity with JumpReLU Sparse Autoencoders 2024.

83. Rajamanoharan, S.; Conmy, A.; Smith, L.; Lieberum, T.; Varma, V.; Kramár, J.; Shah, R.; Nanda, N. Improving Dictionary Learning with Gated Sparse Autoencoders 2024.

84. Lieberum, T.; Rajamanoharan, S.; Conmy, A.; Smith, L.; Sonnerat, N.; Varma, V.; Kramár, J.; Dragan, A.; Shah, R.; Nanda, N. Gemma Scope: Open Sparse Autoencoders Everywhere All at Once on Gemma 2. In Proceedings of the Proceedings of the 7th BlackboxNLP Workshop: Analyzing and Interpreting Neural Networks for NLP; 2024; pp. 278–300.

85. Bussmann, B.; Leask, P.; Nanda, N. BatchTopK Sparse Autoencoders 2024.

86. Bussmann, B.; Nabeshima, N.; Karvonen, A.; Nanda, N. Learning Multi-Level Features with Matryoshka Sparse Autoencoders 2025.

87. Parsan, N.; Yang, D.J.; Yang, J.J. Towards Interpretable Protein Structure Prediction with Sparse Autoencoders 2025.

88. Dunefsky, J.; Chlenski, P.; Nanda, N. Transcoders Find Interpretable Llm Feature Circuits. Advances in Neural Information Processing Systems 2024, 37, 24410–.

89. Wu, W.; Song, X.; Wen, Y.; Lin, Q.; Zhou, Z.; Hu, J.Y.-C.; Wang, Z.; Liu, H. Genome-Factory: An Integrated Library for Tuning, Deploying, and Interpreting Genomic Models 2025.

90. Paulo, G.; Belrose, N. Sparse Autoencoders Trained on the Same Data Learn Different Features 2025.

91. Chanin, D.; Wilken-Smith, J.; Dulka, T.; Bhatnagar, H.; Golechha, S.; Bloom, J. A Is for Absorption: Studying Feature Splitting and Absorption in Sparse Autoencoders 2025.

92. Leask, P.; Bussmann, B.; Pearce, M.; Bloom, J.; Tigges, C.; Moubayed, N.A.; Sharkey, L.; Nanda, N. Sparse Autoencoders Do Not Find Canonical Units of Analysis 2025.

93. Heap, T.; Lawson, T.; Farnik, L.; Aitchison, L. Automated Interpretability Metrics Do Not Distinguish Trained and Random Transformers 2026.

94. Engels, J.; Riggs, L.; Tegmark, M. Decomposing The Dark Matter of Sparse Autoencoders 2025.

95. Smith, L.; Rajamanoharan, S.; Conmy, A.; McDougall, C.; Kramar, J.; Lieberum, T.; Shah, R.; Nanda, N. Negative Results for Sparse Autoencoders On Downstream Tasks and Deprioritising SAE Research. DeepMind Safety Research Blog Post 2025, 1.

96. Wu, Z.; Arora, A.; Geiger, A.; Wang, Z.; Huang, J.; Jurafsky, D.; Manning, C.D.; Potts, C. AxBench: Steering LLMs? Even Simple Baselines Outperform Sparse Autoencoders 2025.

97. Peng, K.; Movva, R.; Kleinberg, J.; Pierson, E.; Garg, N. Use Sparse Autoencoders to Discover Unknown Concepts, Not to Act on Known Concepts 2025.

98. Nainani, J.; Reimer, B.M.; Watts, C.; Jensen, D.; Green, A.G. Mechanistic Evidence That Motif-Gated Domain Recognition Drives Contact Prediction in Protein Language Models 2025.

99. Mudide, A.; Engels, J.; Michaud, E.J.; Tegmark, M.; Witt, C.S. de Efficient Dictionary Learning with Switch Sparse Autoencoders 2025.

100. Kurochkin, V.; Aksenov, Y.; Laptev, D.; Gavrilov, D.; Balagansky, N. Kronecker Factorization Improves Efficiency and Interpretability of Sparse Autoencoders 2025.

101. Karvonen, A.; Rager, C.; Lin, J.; Tigges, C.; Bloom, J.; Chanin, D.; Lau, Y.-T.; Farrell, E.; McDougall, C.; Ayonrinde, K.; et al. SAEBench: A Comprehensive Benchmark for Sparse Autoencoders in Language Model Interpretability 2025.

102. Kim, B.; Wattenberg, M.; Gilmer, J.; Cai, C.; Wexler, J.; Viegas, F. Interpretability beyond Feature Attribution: Quantitative Testing with Concept Activation Vectors (Tcav). In Proceedings of the International conference on machine learning; PMLR, 2018; pp. 2668–2677.

103. Ismail, A.A.; Oikarinen, T.; Wang, A.; Adebayo, J.; Stanton, S.; Joren, T.; Kleinhenz, J.; Goodman, A.; Bravo, H.C.; Cho, K.; et al. Concept Bottleneck Language Models For Protein Design 2024.

